# Quantitative super-resolution imaging of platelet degranulation reveals differential release of VWF and VWF propeptide from alpha-granules

**DOI:** 10.1101/2022.10.25.513669

**Authors:** Maurice Swinkels, Sophie Hordijk, Petra E. Bürgisser, Johan A. Slotman, Tom Carter, Frank W.G. Leebeek, A.J. Gerard Jansen, Jan Voorberg, Ruben Bierings

## Abstract

**Background:** Platelet alpha-granules contain Von Willebrand factor (VWF), which is stored in eccentric alpha-granule nanodomains, and VWF propeptide (VWFpp). Differential release of VWF and VWFpp has been reported from endothelial cells. It is unclear if this also occurs during platelet alpha-granule exocytosis. We have recently developed a 3D super-resolution imaging workflow for quantification of platelet alpha-granule content based on Structured Illumination Microscopy (SIM). With this we can study alpha-granule cargo release following platelet activation in hundreds of platelets simultaneously.

**Aims:** To study release of VWF and VWFpp from alpha-granules using quantitative super-resolution microscopy.

**Methods:** Platelets were activated with PAR-1 activating peptide (PAR-1 ap) or collagen-related peptide (CRP-XL). Alpha-tubulin, VWF, VWFpp, SPARC and fibrinogen were imaged using 3D-SIM, followed by semi-automated analysis in FIJI. Uptake of anti-VWF nanobody during degranulation was used to identify alpha-granules that partially released content.

**Results:** VWF+ and VWFpp+ structures overlapped nearly completely (∼90%) in resting platelets, implying they are stored in similar eccentric alpha-granule nanodomains. A subset of VWF+/VWFpp+-structures was released completely at 0.6 µM PAR-1 ap, but at higher concentration (20 µM) significantly more VWFpp (85.3±1.6%) was released than VWF (37.6±1.4%). Release of other cargo was intermediate at 20 µM (SPARC: 62.2±1.4%; fibrinogen: 51.9±2.9%), providing further evidence for differential cargo release. Similar results were obtained using CRP-XL. Anti-VWF nanobody was taken up by VWF+/VWFpp-structures and increased with stimulus strength, demonstrating these were post-exocytotic structures.

**Conclusions:** VWF and VWFpp are differentially released from alpha-granules. This may affect how platelet-derived VWF and VWFpp contribute to formation and stabilization of hemostatic clots.

**Key points:**

1. VWFpp and VWF are localized in the same, eccentric alpha-granule subdomain in resting platelets and do not overlap with other alpha-granule cargo proteins such as fibrinogen
2. VWFpp and VWF are differentially secreted from individual alpha-granules upon activation with platelet agonists PAR-1 activating peptide and collagen-related peptide

## Introduction

During thrombopoiesis several types of secretory granules from bone marrow megakaryocytes are packaged into budding platelets. Release of their content enables platelets to rapidly respond to changes in their environment, such as during injury, inflammation or when encountering pathogens. alpha-granules are the most abundant platelet secretory organelle, and contain various proteins and molecules involved in the hemostatic response [1,2]. Among these is Von Willebrand Factor (VWF), a key hemostatic adhesive glycoprotein synthesized by megakaryocytes and endothelial cells, whose two main roles are to facilitate platelet adhesion to sites of vascular injury and to stabilize coagulation factor VIII in the circulation [3]. VWF is stored in platelet alpha-granules and in endothelial cell Weibel-Palade bodies (WPBs) and can, together with other granule cargo proteins, be released via exocytosis following cellular activation. Circulating VWF levels in plasma are primarily maintained through basal secretion of WPBs from the endothelium [4].

Our knowledge on VWF biosynthesis primarily comes from studies utilizing endothelial cells and heterologous expression systems as cellular models. As it progresses through the secretory pathway, VWF undergoes several post-translational processing steps which include dimerization, glycosylation and multimerization into long platelet-adhesive concatemers [3]. Within the acidifying milieu of the Golgi, VWF multimers condense into alpha-helical VWF tubules that lend the WPBs their characteristic rod-like shape [5]. Here, a large N-terminal moiety called the VWF propeptide (VWFpp), is also proteolytically cleaved from the mature VWF chain. In endothelial cells, cleaved VWFpp remains non-covalently associated with VWF due to the prevailing conditions in the Golgi and beyond (low pH, high Ca^2+^), leading to its co-packaging in the forming WPBs [6]. VWFpp is essential for VWF multimerization, tubulation and WPB biogenesis [7–9] and becomes an integral part of the VWF tubules *in vitro* and *in vivo* [10,11]. During exocytosis, the vesicle interior neutralizes leading to the rapid decondensation of VWF tubules [12,13] and loss of the non-covalent association between VWF and VWFpp [14]. Depending on the type of exocytosis (full fusion, lingering kiss or compound fusion) [4] and the extracellular environment, VWF and VWFpp undergo divergent fates post-release [14–17].

In platelets, VWF is zonally packaged within eccentric alpha-granule nanodomains which also contain short VWF tubules [18–20]. Platelets also contain VWFpp [21] and are able to secrete it following stimulation with various agonists that induce alpha-granule release [22]. However, the organization of VWFpp in alpha-granules or its release from alpha-granules have not been documented in detail. Similar to endothelial WPBs, platelet alpha-granules can undergo single and compound exocytosis depending on the type and magnitude of stimulus [23,24] and can selectively release small compounds such as chemokines versus larger proteins [25,26]. It is not clear how these processes influence the efficiency of release of VWF and VWFpp specifically, and whether or not VWF and VWFpp release from platelet alpha-granules is comparable to their release from endothelial cell storage organelles.

In this study we have investigated the storage and release of VWF and VWFpp in platelets using 3D Structured Illumination Microscopy (3D-SIM). We show that VWF and VWFpp reside in a distinct alpha-granule subdomain not occupied by other alpha-granule proteins such as fibrinogen. By quantitative 3D-SIM analysis of residual VWF and VWFpp in activated platelets we demonstrate that VWFpp is efficiently released from platelets in a dose-dependent manner, while even at maximal activation the bulk of VWF remains associated with platelets in post-fusion structures. Our study sheds new light on the divergent outcomes of VWF and VWFpp following release from platelet alpha-granules.

## Methods

### Platelet isolation

All steps are carried out at room temperature (RT) unless otherwise stated. Whole blood is drawn from consenting healthy donors in citrate tubes. Washed platelets were prepared as described previously [20]. In brief, platelet-rich plasma (PRP) is generated by centrifugation at 120 x g for 20 minutes with low acceleration (max. 5) and low brake (max. 3). PRP is washed once in 10% acid-citrate dextrose buffer (85 mM Na_3_-citrate, 71 mM citric acid, 111 mM glucose) with 111 µM prostaglandin E_1_ (Sigma), twice in washing buffer (36 mM citric acid, 103 mM NaCl, 5 mM KCl, 5 mM EDTA, 5.6 mM glucose, pH 6.5) with 11 and 0 µM prostaglandin E_1_ respectively, then resuspended at 250*10^3^ platelets/µL in assay buffer (10 mM HEPES, 140 mM NaCl, 3 mM KCl, 0.5 mM MgCl_2_, 10 mM glucose and 0.5 mM NaHCO_3_, pH 7.4).

### Platelet activation

Washed platelets at 250*10^3^ platelets/µL were stimulated with 0-20 µM of PAR-1 activating peptide (Peptides International) or 0-1 µg/ml collagen-related peptide (CRP-XL, CambCol Labs) for 30 minutes at 37 ºC. Reactions are stopped by adding 1% paraformaldehyde (final concentration) for 5 minutes, then quenched with 50 mM NH_4_Cl for 5 minutes. Samples are diluted in a large volume of washing buffer, washed once, and resuspended in assay buffer at approximately 250*10^3^/µL.

### VWF nanobody internalization assay

Washed platelets were incubated with nanobodies directed against the VWF CTCK domain or control nanobodies (s-VWF and R2, respectively [27]; kindly supplied by Dr. Coen Maas, UMCU, Netherlands) at a final concentration of 1 µg/ml and were stimulated as described above. Internalized nanobodies were detected using goat anti-Alpaca IgG-AF488 (Jackson ImmunoResearch).

### Flow cytometry

Small aliquots are taken for quality control of platelet activation by flow cytometry. Samples are stained with CD61-APC (BD Biosciences, 1:400) and CD62P-PE (BD Biosciences, 1:100) or with secondary anti-Alpaca IgG-AF488 (Jackson ImmunoResearch, 1:400) for 15 minutes at RT, diluted in assay buffer, and immediately read on a FACS Canto II flow cytometer (BD Biosciences). In some cases fixed platelets were permeabilized with 0.05% saponin before staining. FSC and SSC parameters were used to gate platelets and single cells, while single stains and isotype controls were used to determine fluorescence gating.

### Platelet seeding and immunofluorescence

Seeding and staining was performed as described previously [20]. In brief, all unique sample conditions were seeded on poly-D-lysine coated 9 mm diameter 1.5H high-precision coverslips (Marienfeld), permeabilized, and stored in PGAS (0.2% gelatin, 0.02% azide and 0.02% saponin in PBS). Primary and secondary antibody staining were done in PGAS for 30 minutes at RT, washed 3x with PGAS following incubations. Antibodies used are listed in Supplementary Table S1. Finally, slides were dipped in PBS, mounted in Mowiol and imaged within one week.

### Structured illumination- and confocal microscopy and image analysis

All samples were imaged with SIM (Elyra PS.1, Zeiss) and confocal microscopy (SP8, Leica). Nanobody internalization assay SIM images were all reconstructed with an identical reference image parallel in Z in order to control for contrast-stretching by the reconstruction algorithm. Due to very bright alpha-tubulin signals and relatively broad emission filters crosstalk between the far-red and red channel was observed. Crosstalk between the channels. This was corrected equally in all applicable images by subtracting the far-red channel (alpha-tubulin) from the red channel. SIM images were analyzed through ImageJ-based processing workflows as described in detail previously [20]. In brief, individual platelets were segmented based on alpha-tubulin staining, and individual 3D granular structures were quantified based on individual staining (e.g. VWF/VWFpp) by automated thresholding. Qualitative images without alpha-tubulin were manually segmented in individual platelets, and were scored for VWF/VWFpp/nanobody overlap by three individual researchers by assessing VWFpp and nanobody positivity in VWF-positive granules defined by our regular ImageJ workflow.

### Immunoblotting

HUVECs (grown as previously described [28]) and washed platelets were lysed in NP-40 buffer (0.5% NP-40, 150 mM NaCl, 10 mM Tris, 5 mM EDTA, pH 8.5). Lysates were separated on 4-12% Bis-Tris NuPAGE gels (Invitrogen) under reducing conditions and transferred to 0.2 µm nitrocellulose membranes. Membranes were probed with rabbit anti-VWF (DAKO) and rabbit anti-VWFpp [17] followed by LT680-labeled donkey anti-rabbit secondary antibodies (Li-COR). Membranes were scanned on an Odyssey scanner (Li-COR).

### Statistical analysis

Individual stimulation conditions were compared with resting platelets by two-way ANOVA. Multiple comparisons were corrected using Sidak’s multiple comparisons test. Dose-response relationships were assessed using linear regression and reported as IC_50_ values. All statistical analyses were performed with GraphPad Prism (version 8). Data is presented as mean ± standard deviation. A p-value under 0.05 was considered statistically significant.

## Results

### VWF propeptide colocalizes with mature VWF in eccentric alpha-granule nanodomains

The localization of VWF and VWFpp in resting platelets was studied by 3D-SIM [20]. VWF is found in platelet alpha-granules (Figure 1A) and in keeping with several ultrastructural studies [18–20,29] we observed discrete volumes of VWF in eccentrically located subdomains within the encapsulation of a P-selectin positive alpha-granule membrane (Supplemental Figure 1). As observed previously [20,30,31], only limited overlap exists with other alpha-granule constituents like fibrinogen and SPARC, which are more diffusely localized inside the alpha-granule matrix (Figure 1B-C) and appear to be excluded from these VWF containing nanodomains. VWFpp is also found in alpha -granules and shows a high degree of overlap with VWF (Figure 1A). We find that VWFpp is also encapsulated by a P-selectin containing membrane and is similarly found in an eccentric position within the alpha-granule colocalizing with VWF (Supplemental Figure 1), suggesting they both occupy the same alpha-granule subdomain. For this analysis we used a polyclonal antibody directed against an octapeptide in the exposed carboxyterminal end of VWFpp following proteolytic cleavage from the mature VWF chain [17]. Immunoblot analysis showed that in platelet lysates this VWFpp antibody exclusively recognizes a 100 kDa protein corresponding to the size of VWFpp (Supplemental Figure 2). Furthermore, contrary to endothelial cells, platelets only contain mature VWF and no detectable proVWF, which suggests that proteolytic processing of proVWF into mature VWF and VWFpp is completed before or during the formation of alpha-granules in megakaryocytes and does not continue post-budding of platelets (Supplemental Figure 2). Together, this further emphasizes that the nearly perfect overlap of VWF and VWFpp in our SIM analysis is likely the result of both being in the same supramolecular structures within alpha-granules and not of cross-reaction of the VWFpp antibody with unprocessed proVWF.

**Figure 1:**
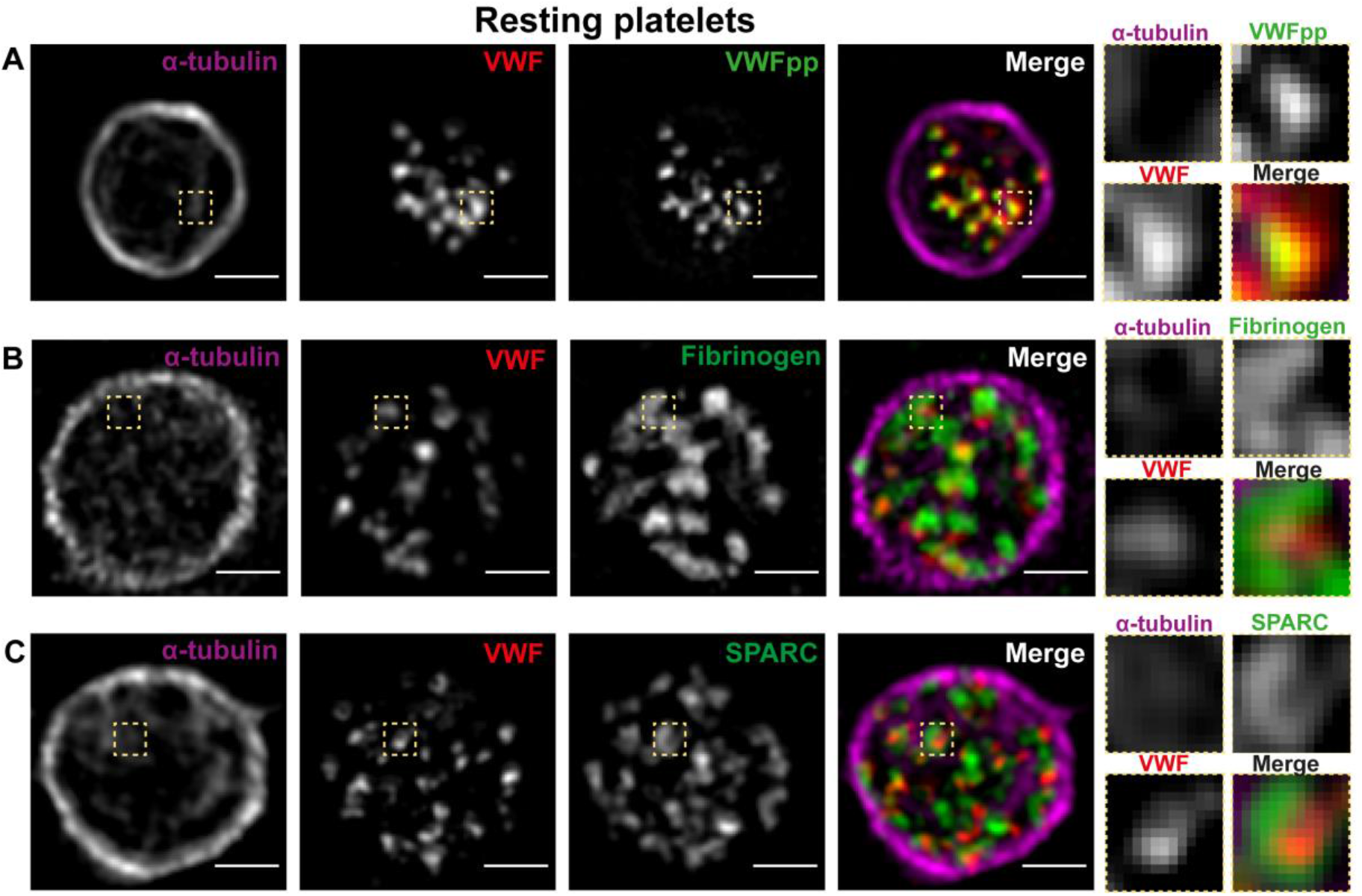
VWF and VWFpp localization in resting platelets. Resting platelets were stained for alpha-tubulin (magenta), VWF (red) and VWFpp (A, green), Fibrinogen (B) or SPARC (C). Imaging was done by SIM and high-resolution single Z-slices are shown of representative images, including zoom-ins (yellow square). Scale bar represents 1 µm.

### Platelet stimulation triggers differential release of VWFpp and VWF

We then investigated VWF and VWFpp secretion from platelet alpha-granules following platelet activation induced by PAR-1 and GPVI signaling. We first evaluated single VWF- and VWFpp-granule release by quantifying remaining VWF+ and VWFpp+ structures by 3D-SIM (Figure 2). We observed only limited VWF release from platelets after PAR-1 stimulation (20 µM PAR-1 ap) (Figure 2A), with VWF often remaining in granular structures or in some cases within larger, centrally located clusters (Figure 2B). This incomplete release of VWF has been previously documented and is not due to suboptimal stimulation, as also evidenced by the high proportion of degranulating platelets (Supplemental Figure 3). Interestingly, we found that residual VWF staining is confined to P-selectin (CD62P) labeled structures, suggesting it mostly remains in (post-exocytotic) alpha-granules (Supplemental Figure 4). In contrast, we found that there was profound VWFpp release from alpha-granules under these conditions, illustrated by the loss of VWFpp+ structures and the loss of VWFpp+ immunoreactivity in the remaining VWF containing structures (Figure 2A). Additional examples illustrate there was some variation across platelets (Figure 2B), but the overall effect pivots to clear differential release from alpha-granules between VWF and VWFpp. Similar differences between remaining VWF and VWFpp was observed after stimulation with 1 µg/ml CRP-XL, which triggers the GPVI pathway (Figure 2C). Together, this shows that VWF and VWFpp, despite their close proximity within alpha-granules in resting platelets, can be differentially released by activated platelets. As differences in VWF and VWFpp release in relation to agonist responsiveness may be explained by the large differences in size between VWF and VWFpp (VWFpp is a 100 kDa protein, while ultra-large VWF multimers can be in excess of 100,000 kDa), we also looked at exocytosis of other alpha-granule constituents. SPARC (40 kDa) was released slightly more extensively than VWF (Figure 3A), while fibrinogen (∼340 kDa) release appeared more similar to VWF (Figure 3B). This would suggest that additional factors other than protein size play a role in facilitating the differential agonist responsiveness of VWF versus VWFpp.

**Figure 2:**
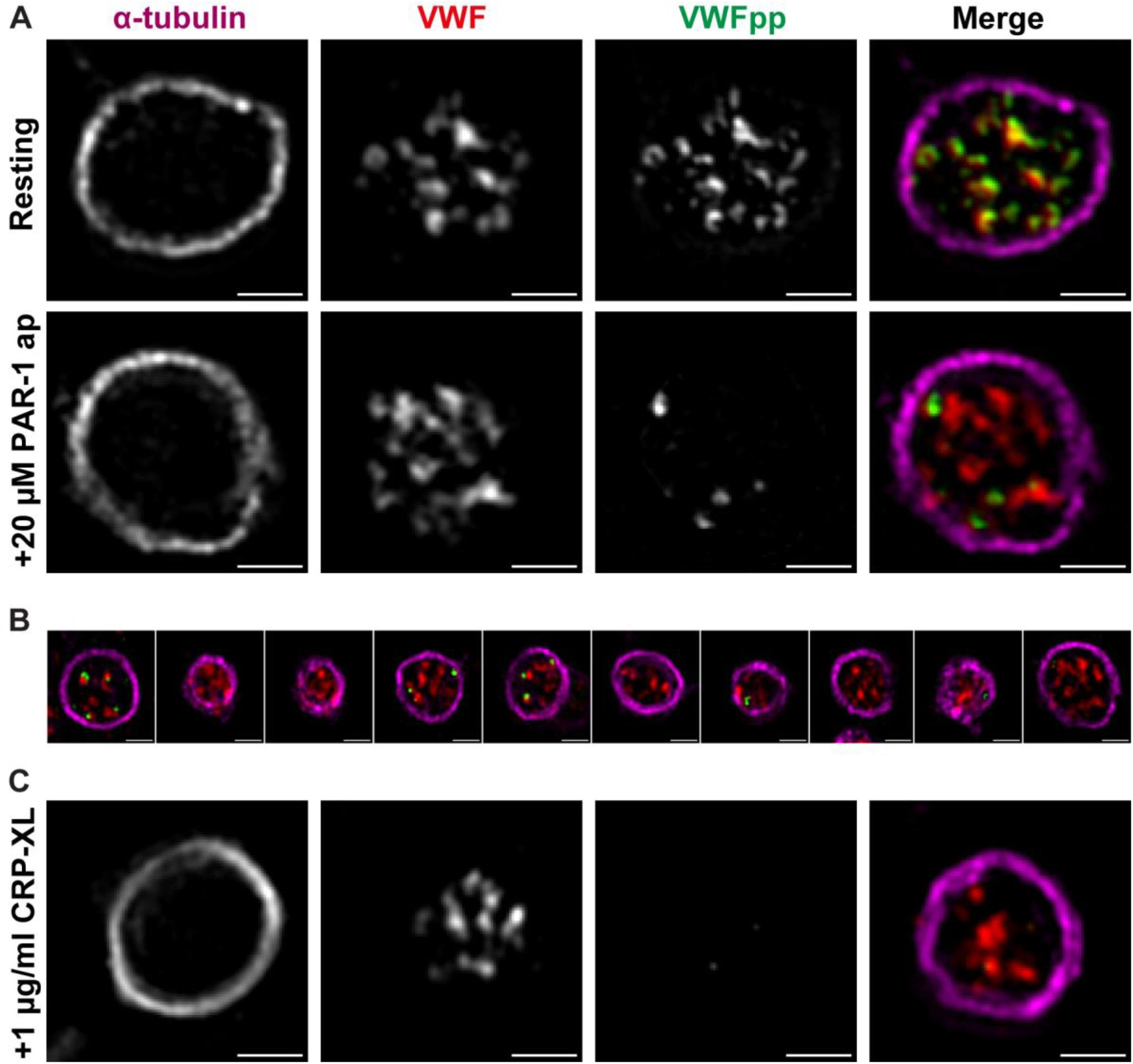
Alpha-granule release of VWF and VWFpp. Platelets were stimulated with 20 µM PAR-1 ap (A-B) or 1 µg/ml CRP (C) and compared to resting platelets for release of VWF and VWFpp. Single plane zoom-in images are shown as well as panels with 10 random platelets. Scale bar represents 1 µm.

**Figure 3:**
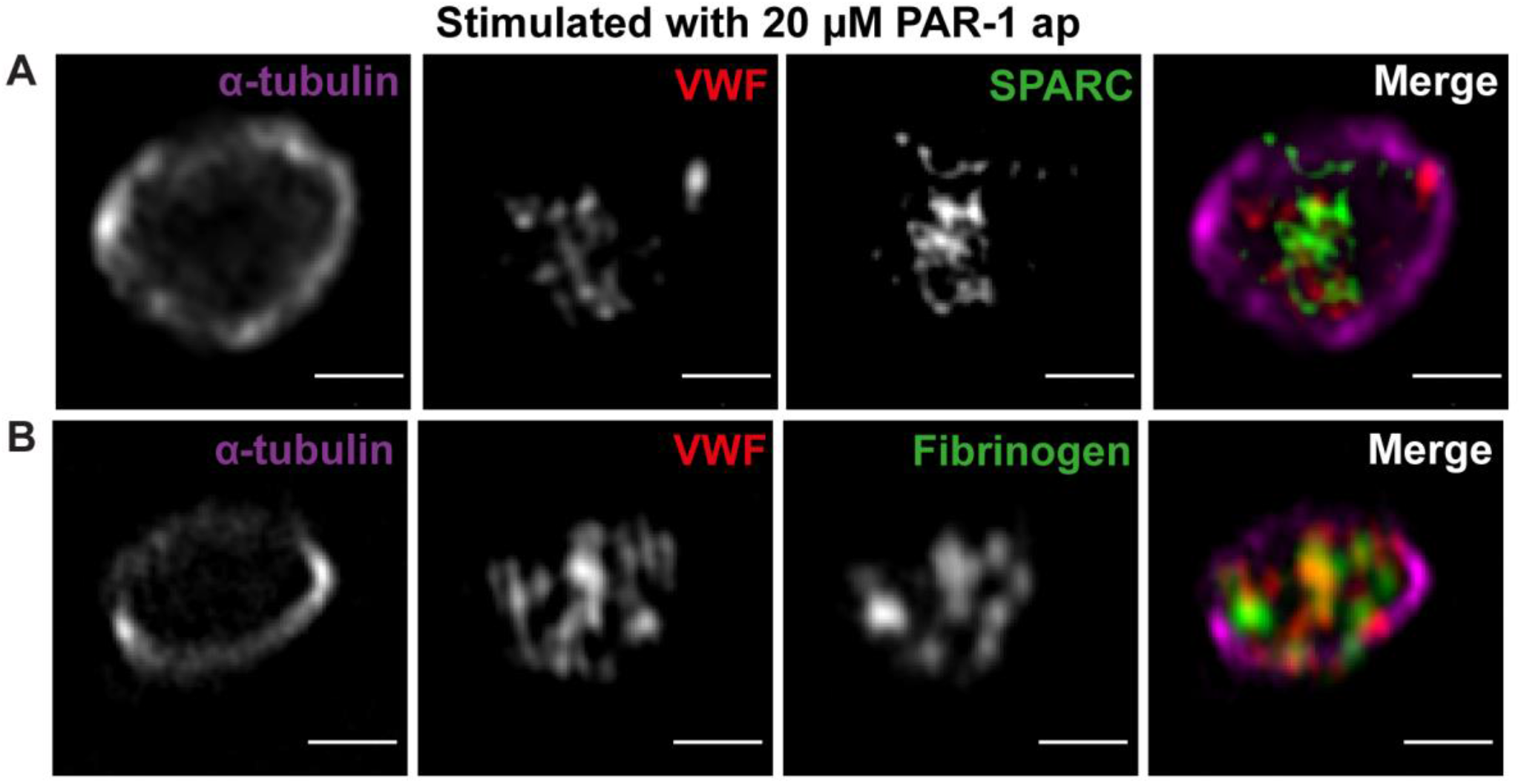
Alpha-granule release of fibrinogen and SPARC. Platelets were stimulated with 20 µM PAR-1 and compared to resting platelets for release of VWF and fibrinogen (A) or SPARC (B). Single plane, representative zoom-in images are shown. Scale bar represents 1 µm.

### Differential release of VWF and VWFpp relates to agonist responsiveness

Next, we investigated whether the differential release of VWF and VWFpp was dependent on agonist strength by using a semi-automated quantitative workflow on 3D-SIM images [20] of platelets activated with a broad concentration range of PAR-1 and CRP-XL. Using additional agonist concentrations that partially or fully trigger alpha-granule release (Supplemental Figure 3), we found that differential release of VWF and VWFpp was apparent at all stimulus concentrations of (Figure 4A), similar to CRP-XL (Supplemental Figure 5). Compared to resting platelets, more VWFpp was released from platelets stimulated with the lowest agonist concentration (0.625 µM PAR-1 ap) (**73.1%** residual VWFpp granules, **p=0.06**) and this was increasingly prominent at 2.5 µM (**42.8%, p=0.0003**) and 20 µM (**21.2%, p<0.0001**). In contrast, VWF release was not significant at any concentration compared to resting platelets, whereas maximal release of VWF at 20 µM (**75.6%** residual VWF granules) was still lower than VWFpp release at 0.625 µM. Indeed, dose-response curves of PAR-1-mediated VWFpp- and VWF-release were significantly different as determined by regression analysis (IC_50_ = **2.01** (VWFpp) vs. IC_50_ = **56.70** (VWF), **p<0.0001**). Similarly, dose-response curves of CRP-XL mediated release were different (IC_50_ = **1.09** (VWFpp) vs. IC_50_ > **100** (VWF), **p<0.0001)**.

**Figure 4:**
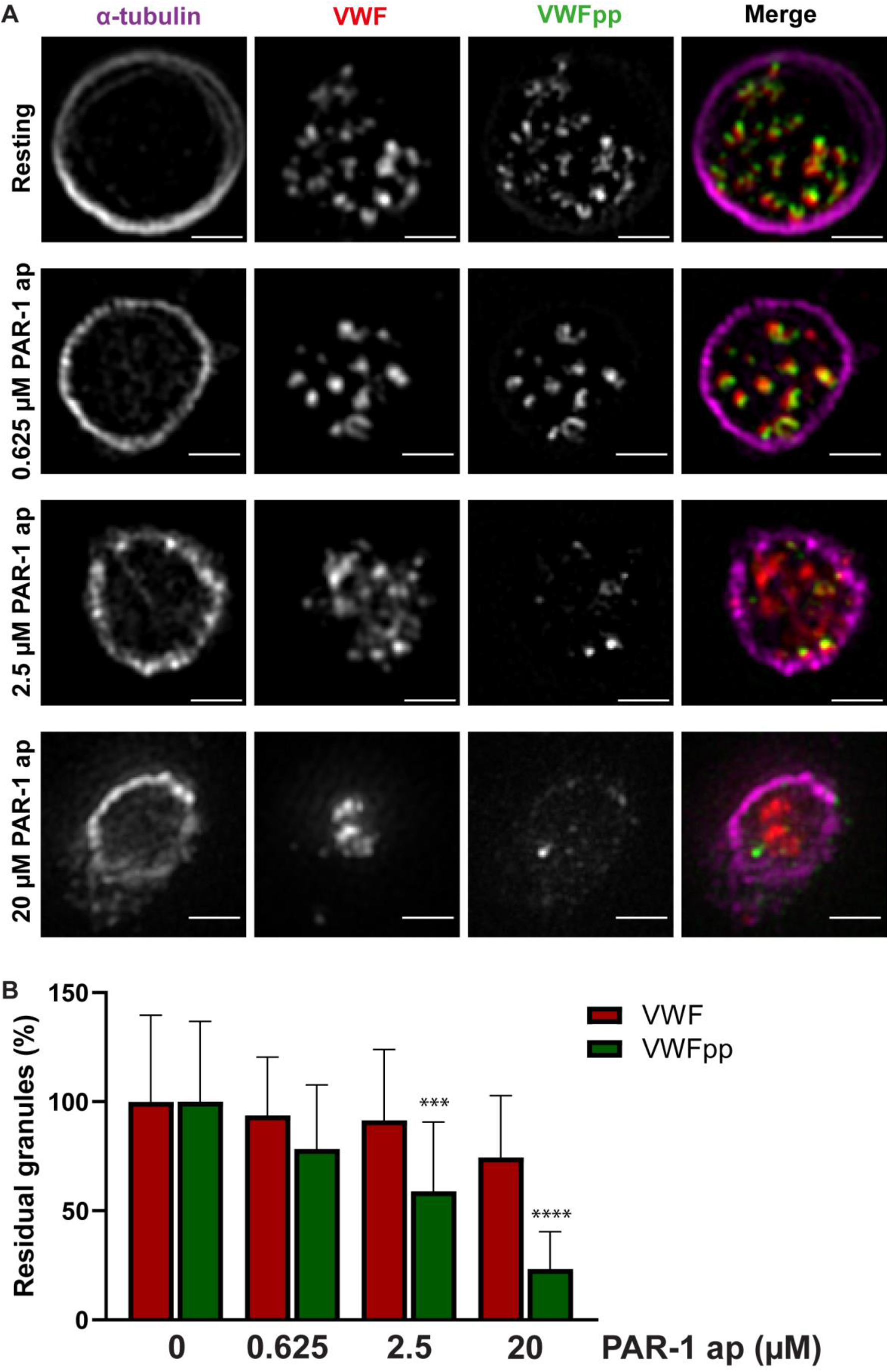
Dose-response release of VWF and VWFpp. Platelets were stimulated with 0-20 µM PAR-1 ap and stained for alpha-tubulin, VWF and VWFpp (A). Representative single plane zoom-in images are shown. Scale bar represents 1 µm. VWF and VWFpp release were assessed by quantification of their residual levels in platelets normalized to resting platelets (B). *** = p<0.001, **** = p<0.0001. Error bars represent SEM / SD.

In contrast, dose-response curves of SPARC and fibrinogen release did not differ from VWF (**p=0.23** (SPARC vs. VWF) and **p=0.31** (fibrinogen vs. VWF)), illustrating differential release of VWFpp and VWF is specific for these two proteins. The intermediate release of these proteins illustrates that the differential release of VWF versus VWFpp is not a constant for different proteins.

In conclusion, we observe a large disparity in alpha-granule release of VWF versus VWFpp, where the former is partially retained in alpha-granules, even under strong stimulatory conditions. In contrast, VWFpp release is sensitive to lower agonist concentrations of PAR-1.

### Anti-VWF nanobody incorporates in post-exocytotic VWF^+^ structures in degranulation-dependent manner

Finally, we wanted to study how and when individual alpha-granular structures differentially release VWF versus VWFpp. As we clearly identified granule populations that contained residual VWF, but no more VWFpp, this would suggest that individual alpha-granules could perform a kiss-and-run type of exocytosis that facilitates release of selective alpha-granule cargo.

To investigate this further, we performed a platelet degranulation experiment with an anti-VWF nanobody (s-VWF) added in suspension, under the assumption that opening of an alpha-granule during exocytosis would facilitate uptake of the nanobody. We found that uptake of the nanobody was directly dependent on the degree of platelet stimulation and thus degranulation, whereas a control R2 nanobody non-specific for VWF did not show any signal by flow cytometry (Figure 5A). Additionally, permeabilised platelets showed an increasingly higher MFI at higher doses of PAR-1, suggesting increasing amounts of nanobody specifically inside platelets (Figure 5A). We further confirmed this with confocal imaging, where we observed accumulation of the nanobody inside the tubulin-ring at 20 µM PAR-1 ap but not in resting platelets (Figure 5B). Additionally, the nanobody co-localized completely with residual VWF^+^-structures suggesting that all VWF^+^ granules are post-exocytotic under these conditions. Together, these findings show that uptake of the VWF nanobody is degranulation dependent.

**Figure 5:**
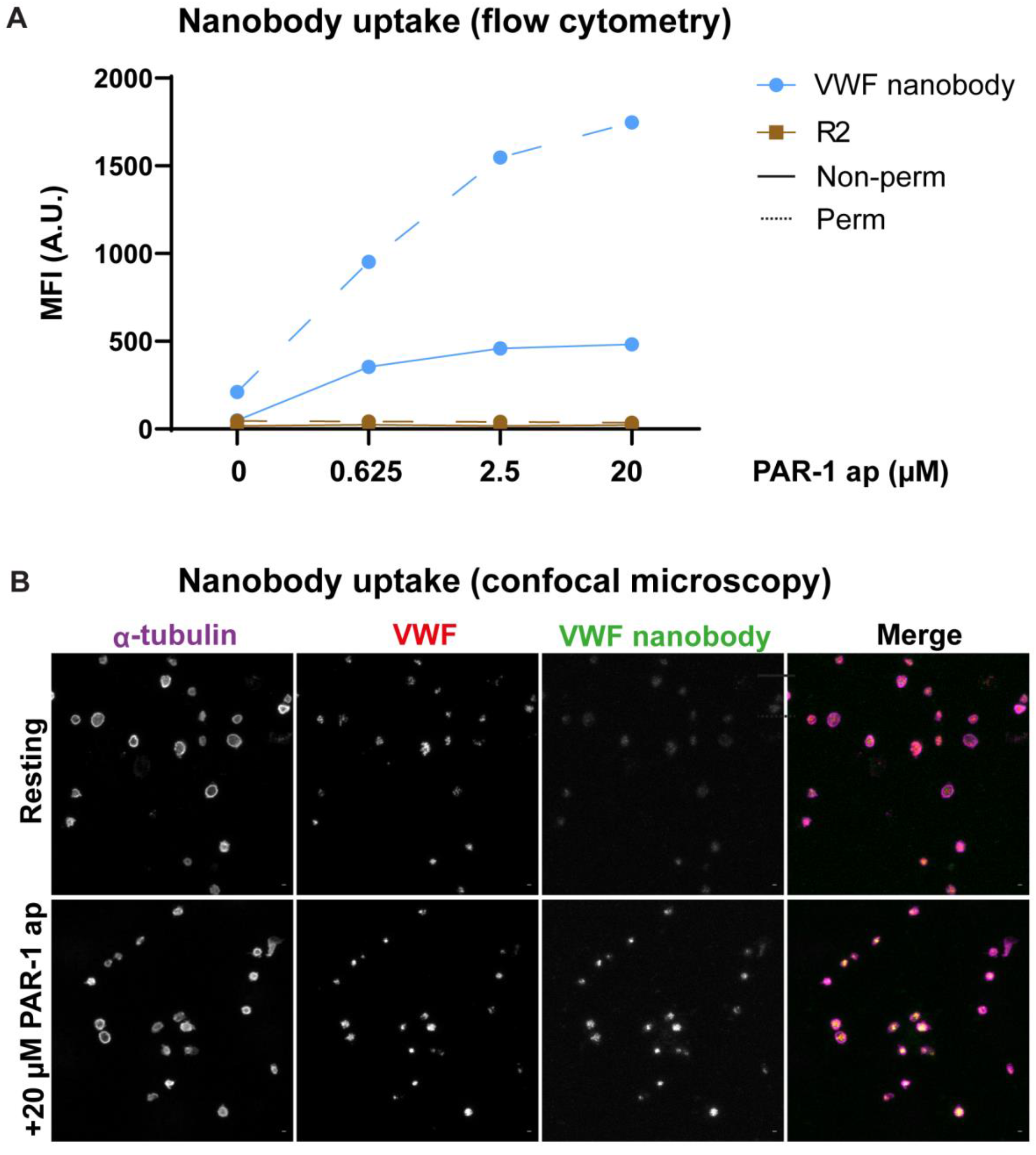
Nanobody bulk uptake in platelets. Stimulation of platelets with 0-20 µM PAR-1 in presence of 1 µg/ml VWF nanobody or R2 control nanobody was analyzed for nanobody uptake by flow cytometry (A) and confocal microscopy (B).

Ultimately, we analyzed individual alpha-granules that were able to take up the VWF nanobody through 3D-SIM. In accordance with the flow cytometry and confocal data, we found an increasing population of VWF nanobody^+^ structures co-localizing with residual VWF that was directly related to the degree of stimulation (Fig 6). At a low dose of PAR-1 ap (Fig 6A), only a minority of granules was strongly positive for the nanobody. These were also VWF^+^ and VWFpp^+^ (zoom-in, Fig 6A). The majority of granules however was VWF^+^ and VWFpp^+^ but revealed weakly staining for the nanobody.

**Figure 6:**
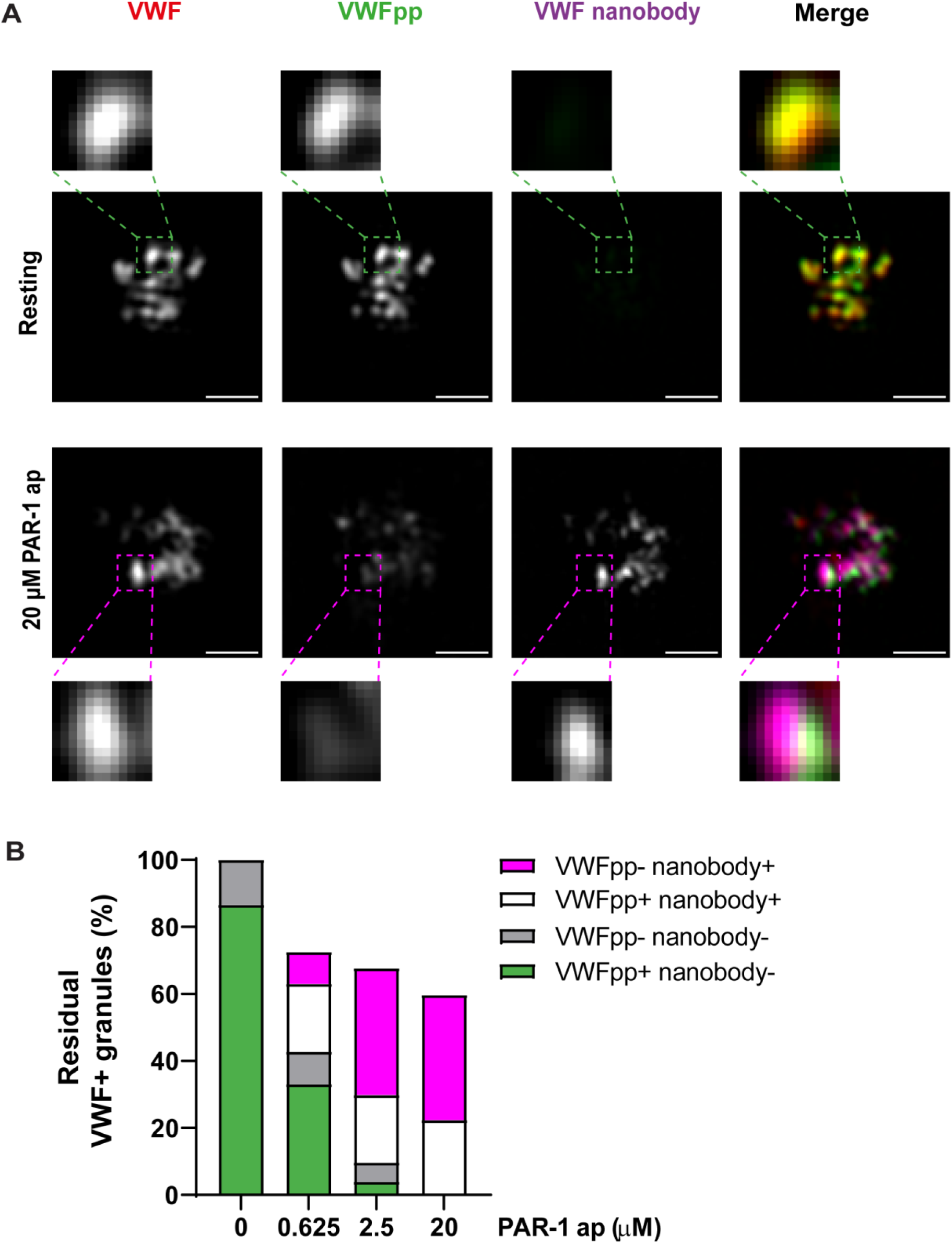
Nanobody uptake analysis in high resolution. Platelets were stimulated with varying doses of PAR-1 ap (A) and granule populations of VWF/VWFpp/VWF-nanobody positivity were assessed (B). Scale bar represents 1 µm.

At a higher dose of PAR-1 ap (Fig 6B), we found a majority of VWF nanobody^+^ and VWF^+^ granules, but these did not contain any VWFpp (zoom-in, Fig 6B), suggesting this content has been released during granule opening. Taken together, our findings imply that increasing doses of PAR-1 ap trigger large-scale release of VWFpp from alpha-granules, while VWF is partially retained in such post-exocytotic granules as evidenced by PAR-1 dependent accumulation of VWF nanobody in VWFpp-VWF^+^ structures.

Our cumulative findings show that alpha-granules may exclusively release content like VWFpp while maintaining other cargo, like VWF, under the conditions described in our work.

## Discussion

Important biochemical and functional differences exist between platelet and endothelial (plasma) VWF [32] that suggest dissimilarities in biosynthesis of VWF between endothelial cells and megakaryocytes: platelet VWF is composed of higher molecular weight multimers, carries different N-linked glycan structures which makes it more resistant to proteolysis by ADAMTS13 [33] and has higher binding affinity for alphaIIbβ3-integrin [34]. In this study, we investigated storage and exocytosis of VWF and VWFpp from platelet alpha-granules through quantitative super resolution microscopy. Our results show that VWFpp is eccentrically localized within alpha-granules in close proximity to mature VWF. In endothelial cells VWFpp integrates in tubules composed of helically condensed VWF multimers that are found within WPBs. Given that similar tubules, albeit shorter in length, have been observed in platelet alpha-granules [19], we speculate that VWFpp is similarly arranged within VWF tubules as in endothelial WPBs.

But contrary to WPBs, where the tubular arrangement of VWF is essential for rapid and efficient release of VWF upon exocytosis, alpha-granules only release a limited amount of their VWF, even at agonist concentrations that elicit maximum surface exposure of P-selectin and lead to incorporation of anti-VWF nanobody into practically all remaining VWF positive structures. The latter is important because it implies that all these granules have undergone a granule fusion event that generated a fusion pore in contact with the extracellular space. Additionally, we found evidence for differential release of VWFpp and VWF, showing that individual alpha-granules can preferentially release their VWFpp cargo while retaining VWF. Differential release was dependent on agonist responsiveness but not related to the type of agonist we used in our study.

This is in sharp contrast with the 1:1 stoichiometry between VWF and VWFpp that is released from endothelial cells [14]. What could explain the difference in secretion efficiency between VWF and VWFpp from alpha-granules? Earlier studies on the organization and exocytosis of different types of alpha-granule cargo have resulted in several models as to how platelets are able to (differentially) release their content. Based on limited colocalization of a number of alpha-granule cargo proteins, including VWF and fibrinogen as well as several pro- and anti-angiogenic mediators, it was postulated that subpopulations of alpha-granules exist based on their inclusion of cargo with opposing functions [35,36]. Preferential mobilization of one of these subpopulations by specific agonists would then lead to differential release of distinct functional classes of alpha-granule cargo, giving platelets the opportunity to direct their secretory response in a context-specific manner. However, this hypothesis was significantly challenged by quantitative, high-resolution imaging that showed that alpha-granule cargo is stochastically packaged in alpha-granules, but segregated within subdomains of the granule matrix [29–31]. Kinetic release studies also showed little evidence of specific alpha-granule subpopulations, but instead identified 3 classes of cargo release based on their rate constants (fast, intermediate and slow) in which the alpha-granule cargo distribution is random [25]. Several non-mutually exclusive mechanisms have been proposed that can achieve differential release of cargo from the same granule, including exocytotic fusion mode (compound vs. lingering kiss vs. direct fusion), differences in cargo solubilization or polar release of non-homogenously distributed cargo from one side of the granule [25,37]. The nearly perfect overlap between VWF and VWFpp that we observed in resting platelets (Figure 1-2, S2) suggests both proteins are localized in the same alpha-granules and occupy the same granule subdomains, which rules out that the differences in their release were reflective of granule subpopulations or could have been the result of polar release. Differential release through premature closure of the fusion pore, such as in lingering kiss exocytosis [37], is also unlikely to serve as an explanation since the size of VWFpp (∼100 kDa) would require the fusion pore to fully expand before release. Indeed, we did not find an obvious correlation between releasability and size as SPARC (40 kDa) was less sensitive to low concentration stimulation and achieved lower maximal release than VWFpp (Figure 3).

In line with previous reports by others [24,38], we frequently observed a clustering of VWF positive structures in the central area of activated platelets that were negative for VWFpp, especially at higher agonist concentrations (Figure 4, Supplemental Figure 5). In some cases, a continuous P-selectin staining enveloping several VWF positive structures (Supplemental Figure 4) was present, reminiscent of several alpha-granules that had engaged in compound fusion. Possibly, this exocytotic fusion mode poses no obstacle for VWFpp but does not favor the release of bulky, multimeric cargo such as VWF, for instance by preventing the orderly unfurling of VWF tubules. This may indirectly also relate to differences in solubility of VWF and VWFpp, such as previously observed during loss from the cell surface of endothelial cells following release from WPBs [15]. As a result, VWF remains stuck in post-fusion alpha-granules while VWFpp is efficiently released.

While traces of VWFpp may stick to the D’D3 region of VWF post-release [39], it is likely that after exocytosis its extracellular course is primarily VWF-independent, as attested by the large difference in plasma survival between VWF and VWFpp [40]. However, despite the well-documented pleiotropic roles of VWF, the biological function of extracellular VWFpp is still unclear. Several *in vitro* studies have demonstrated that bovine VWFpp can bind to collagen type I [41,42] and that this interaction can block collagen-induced platelet aggregation [43]. VWFpp also contains an RGD sequence, a motif that can serve as a ligand for a subfamily of integrins that contain α5, α8, αv and αIIb subunits. The VWFpp RGD motif is not strongly conserved between species [44], the integrin receptor for this site has not been identified and its significance remains uncertain as the RGD sequence appears to be unfavorably arranged within the native conformation to support adhesive interactions [43]. Bovine VWFpp can bind alpha4β1- and alpha9β1-integrins, which are expressed on lymphocytes, monocytes and neutrophils, via a sequence within the VWD2 domain that is conserved in humans [45–47]. Another ligand for these integrins, coagulation factor FXIII, has been shown to cross link VWFpp to the extracellular matrix protein laminin [47–49]. Possibly, focused release of VWFpp from degranulating platelets during the initial thrombus formation and incorporation in the adhesive surface via laminin and collagen provides a mechanism to influence the adhesive properties of the exposed extracellular matrix and direct hemostatic and immune responses following vascular injury. Recent reports have emerged that VWFpp can support platelet adhesion to collagen surfaces and enhance thrombus mass in a glycan-dependent manner [50] and that in a murine model of deep vein thrombosis VWFpp incorporates in venous thrombi near regions of active thrombus formation [51].

We have recently shown that PF4 levels in plasma are positively correlated with current severity of bleeding phenotype in VWD type 1 patients [52]. PF4 is a chemokine that is mainly produced by megakaryocytes and stored in platelet alpha-granules, which means that systemic PF4 levels are reflective of platelet degranulation. One possible explanation for the observed association with bleeding severity in this group is that apart from a quantitative deficiency of VWF in plasma, the hemostatic contribution of platelets is impaired by premature release of alpha-granules. This could lead to insufficient delivery of their hemostatic content, such as platelet VWF and other alpha-granule cargo, to sites of vascular injury. A number of studies have focused on the role of platelet derived VWF in hemostasis [53–57]. VWD patients with mild and severe circulating VWF deficiencies who still have residual platelet VWF show a milder clinical phenotype [20,58]. Platelet VWF has also been reported to be important for DDAVP-related amelioration of bleeding times in subgroups of type 1 VWD patients [59]. Together this leads to the notion that release of platelet VWF helps to establish hemostasis in these patients. Our data suggest that following activation the majority of mature VWF actually remains within the platelets, well away from supporting any interactions that can contribute to hemostatic functions of platelets such as adhesion or aggregation. This is in contrast to its proteolytic cleavage product VWFpp, which is efficiently released from platelet alpha-granules following activation and has its own capabilities to interact with components of the extracellular matrix, cellular adhesion receptors and the thrombus. The question thus arises how much of the perceived role of platelet VWF to hemostasis can be attributed to mature VWF and how much (if not more) is actually dependent on VWFpp. More studies that focus on the extracellular role(s) of VWFpp, from endothelial as well as platelet origin, are urgently needed.

## Acknowledgements

We thank Dr. Coen Maas (University Medical Center Utrecht, Netherlands) for generous supply of Alpaca anti-VWF CTCK nanobodies. We thank Titus Lemmens (Maastricht University Medical Center+, Netherlands) for stimulating discussion and experimental assistance. This study has been supported by grants from the Landsteiner Stichting voor Bloedtransfusie Research (LSBR-1707 and LSBR-2005) and an EHA Clinical Research Fellowship (AJG Jansen).

## Author Contributions

MS, SH, PEB, JAS and RB performed experiments and analyzed data. TC and FWGL provided essential reagents and expertise. MS, SH, AJGJ, JV and RB designed the research and wrote the paper. All authors critically revised and approved of the final version of the manuscript.

## Conflict of Interest

F.W.G. Leebeek received research support from CSL Behring, Takeda, uniQure and Sobi and is consultant for uniQure, Biomarin, CSL Behring and Takeda, of which the fees go to the institute. He was a DSMB member for a study sponsored by Roche. A.J.G. Jansen received speaker fees and travel cost payments from 3SBio, Amgen and Novartis, is on the international advisory board at Novartis and received research support from CSL Behring, Principia and Argenx. None of the other authors have conflicts of interest to declare.

