## Supplemental Files for "Quantitative super-resolution imaging of platelet degranulation reveals differential release of VWF and VWF propeptide from alpha-granules"

**Supplementary information**

**Supplementary Table S1: Antibodies used in immunofluorescent stainings.**

| **Antigen** | **Species (Isotype)** | **Label** | **Supplier** | **Cat. Nr.** | **Dilution** |
|---|---|---|---|---|---|
| Von Willebrand factor propeptide | Rabbit | - | Prof. Tom Carter, SGUL | - | 1:500 |
| Von Willebrand factor | Rabbit | - | DAKO | A0082 | 1:500 |
| Von Willebrand factor | Mouse ($IgG_{2b}$) | - | Sanquin | CLB-RAg20 | 1:500 |
| Alpha-tubulin | Mouse ($IgG_{2b}$) | - | Abcam | ab56676 | 1:500 |
| Alpha-tubulin | Mouse ($IgG_1$) | - | Sigma | DM1A | 1:500 |
| SPARC | Mouse ($IgG_1$) | - | SantaCruz | sc-73472 | 1:500 |
| Fibrinogen | Rabbit | - | DAKO | A0080 | 1:500 |
| CD62P | Mouse ($IgG_1$) | - | Bio-Rad | MCA796 | 1:500 |
| Mouse $IgG_1$ | Goat | CF488A | Biotium | 20246 | 1:1000 |
| Mouse $IgG_1$ | Goat | CF568 | Biotium | 20248 | 1:1000 |
| Mouse $IgG_1$ | Goat | CF647 | Biotium | 20252 | 1:1000 |
| Mouse $IgG_{2b}$ | Goat | CF647 | Biotium | 20272 | 1:1000 |
| Rabbit IgG (H+L) | Goat | CF488 | Biotium | 20012 | 1:1000 |
| Rabbit IgG (H+L) | Donkey | AF 568 | ThermoFisher | A11042 | 1:400 |
| Alpaca IgG | Goat | AF 488 | Jackson ImmunoResearch | 128-545-230 | 1:400 |

### Supplemental Figure 1

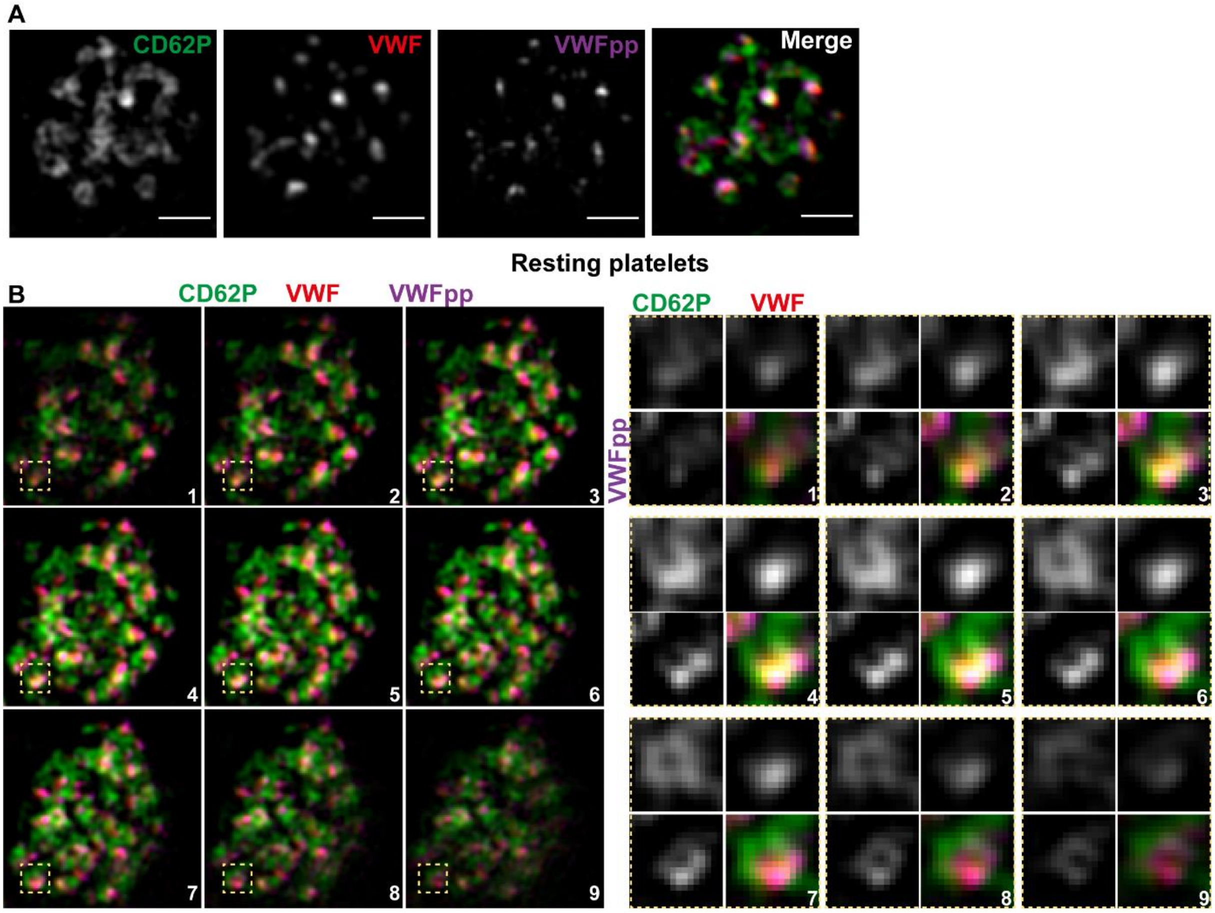


**Supplemental Figure 1: 3D VWF and VWFpp localization in CD62P-defined alpha-granular structures.** Resting platelets were stained for CD62P (green), VWF (red) and VWFpp (magenta). A representative platelet is shown as single plane (A) or in serial sections (B) to illustrate 3D localization of the stained proteins. Single granule zoom-ins of CD62P/VWF/VWFpp (yellow squares) are shown. Scale bar represents 1 µm.

**Supplemental Figure 2**

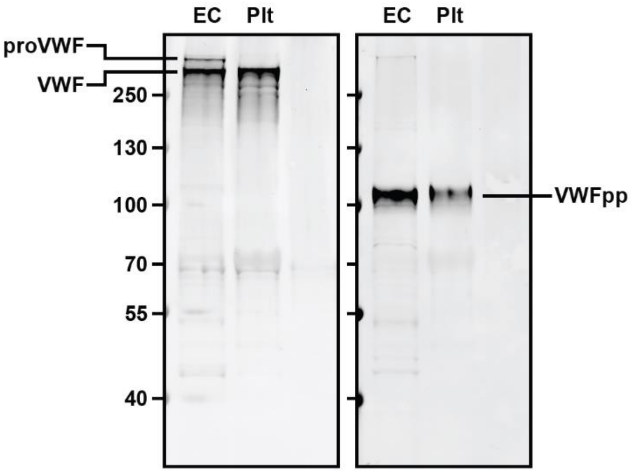


**Supplemental Figure 2: VWF and VWFpp immunoblots of endothelial- and platelet lysates.** Endothelial and platelet lysates were separated on a 4-12% Bis-Tris gel and probed for VWF (left) or VWFpp (right). Bands corresponding to proVWF, mature VWF (VWF) and VWFpp are indicated.

**Supplemental Figure 3**

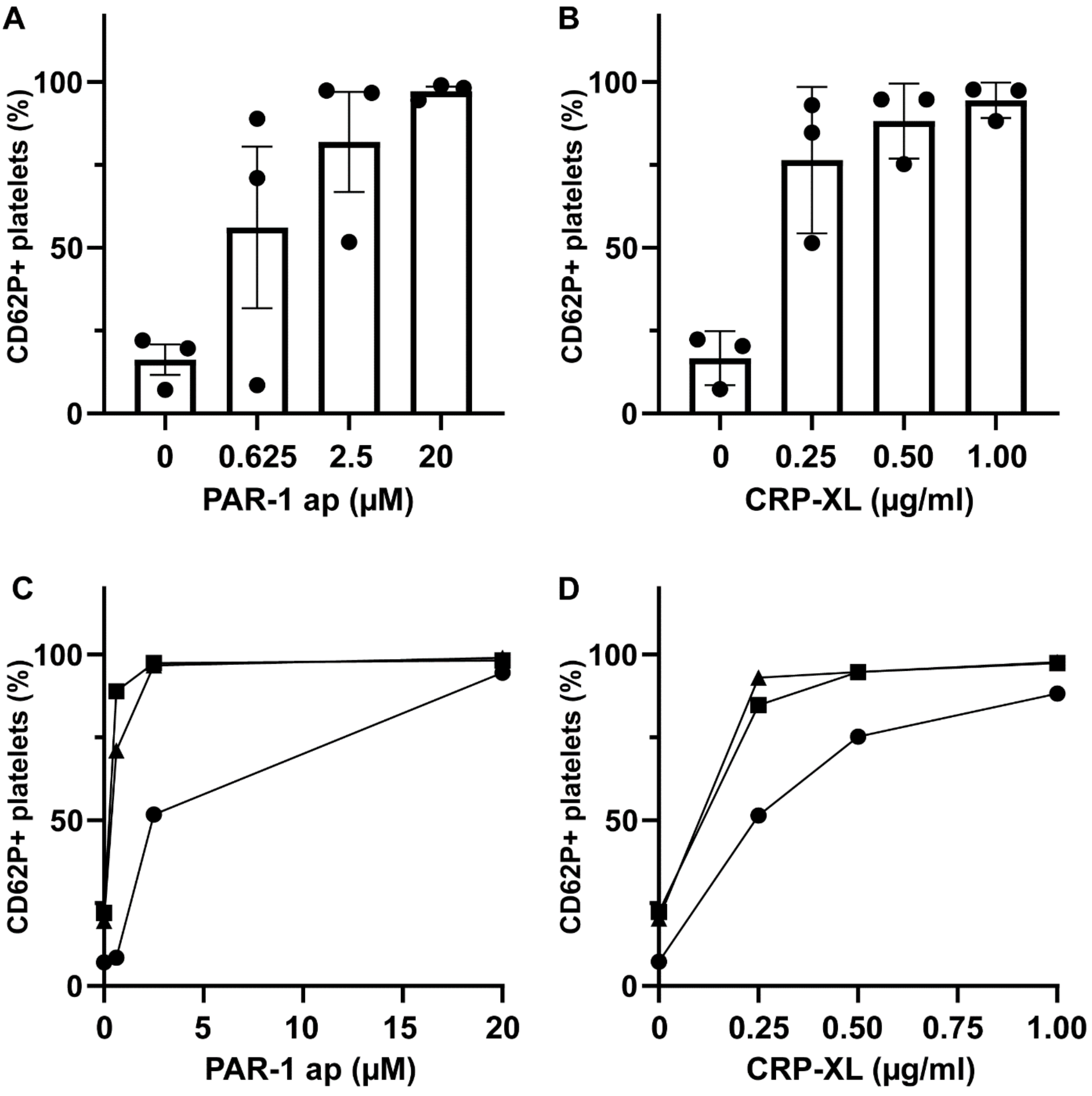


**Supplemental Figure 3: Alpha-granule release assessed by FACS analysis of P-selectin exposure.** Platelets were stimulated with increasing doses of PAR-1 ap (A) or CRP-XL (B) and quantified for CD62P+ cell surface exposure by flow cytometry. Dots represent individual donors (n=3). Individual dose response curves are shown in C (PAR-1 ap) and D (CRP-XL) with symbols representing unique donors. Data presented as mean ± SD.

### Supplemental Figure 4

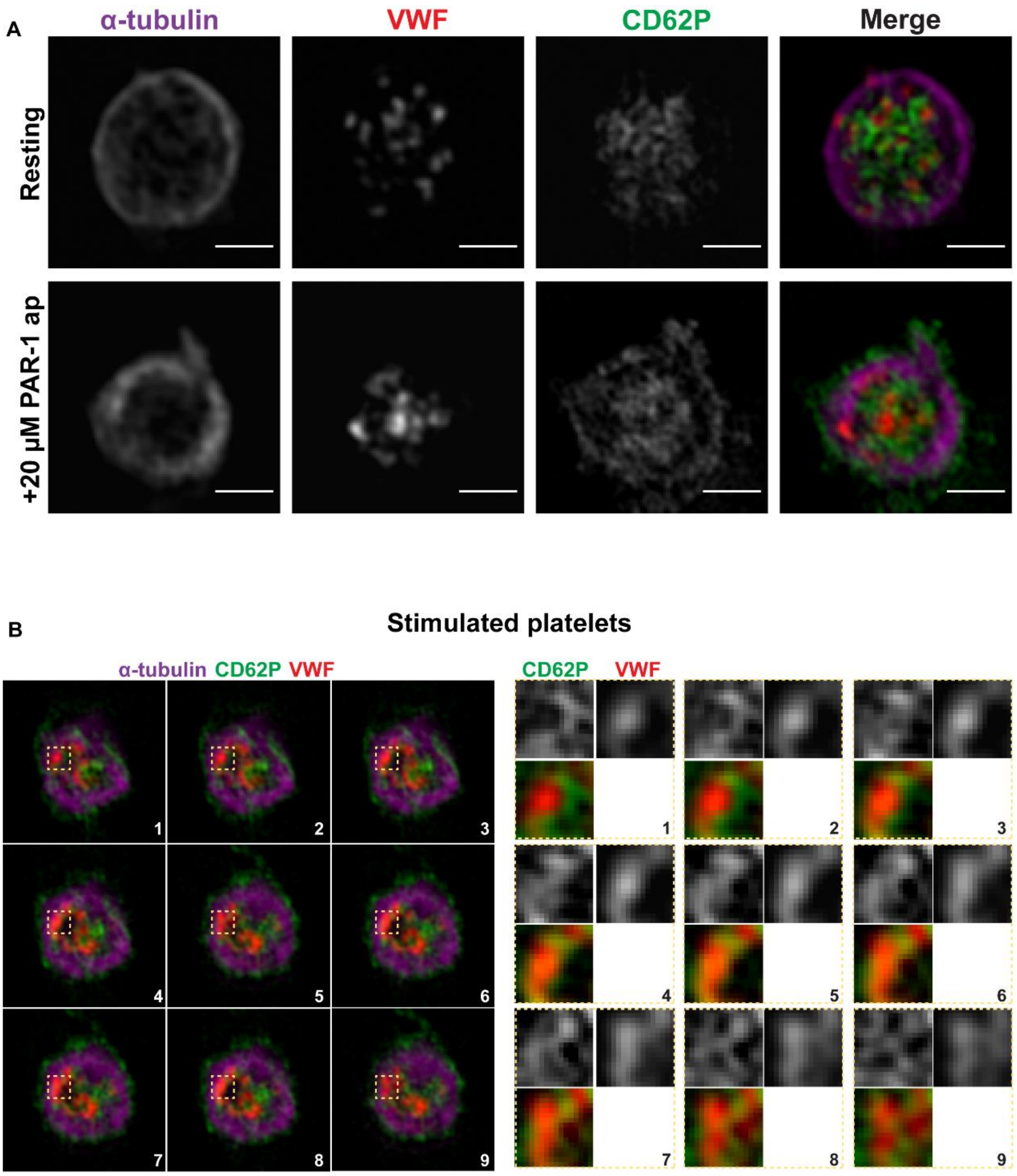


**Supplemental Figure 4: 3D VWF localization in stimulated platelets.** Platelets stimulated with 20 µM PAR-1 ap were stained for CD62P (green), VWF (red) and α-tubulin (magenta) and compared to resting platelets (A). A representative platelet is shown in serial sections and single granule zoom-ins to illustrate 3D localization of CD62P and VWF (B). Scale bar represents 1 µm.

**Supplemental Figure 5**

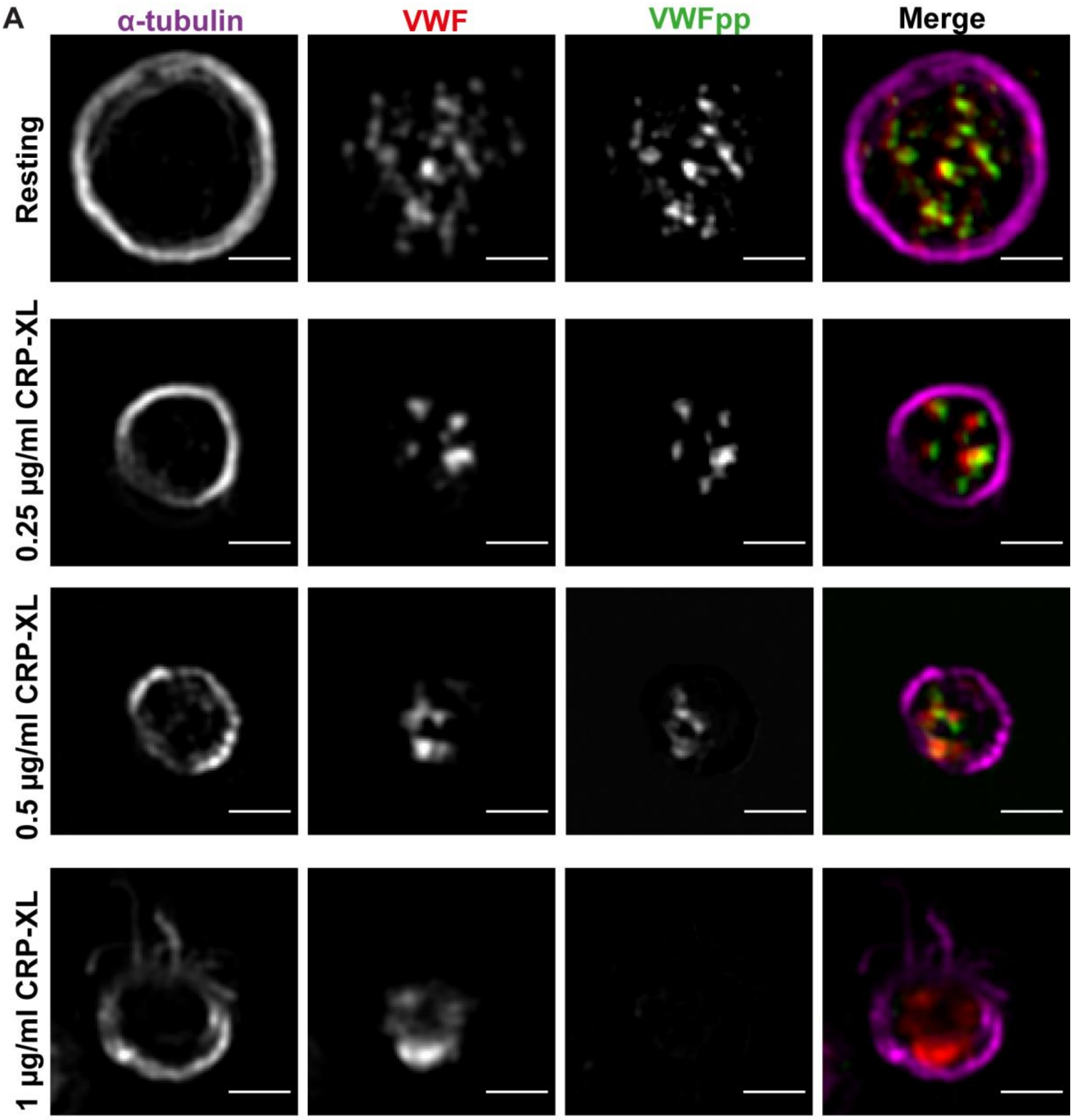


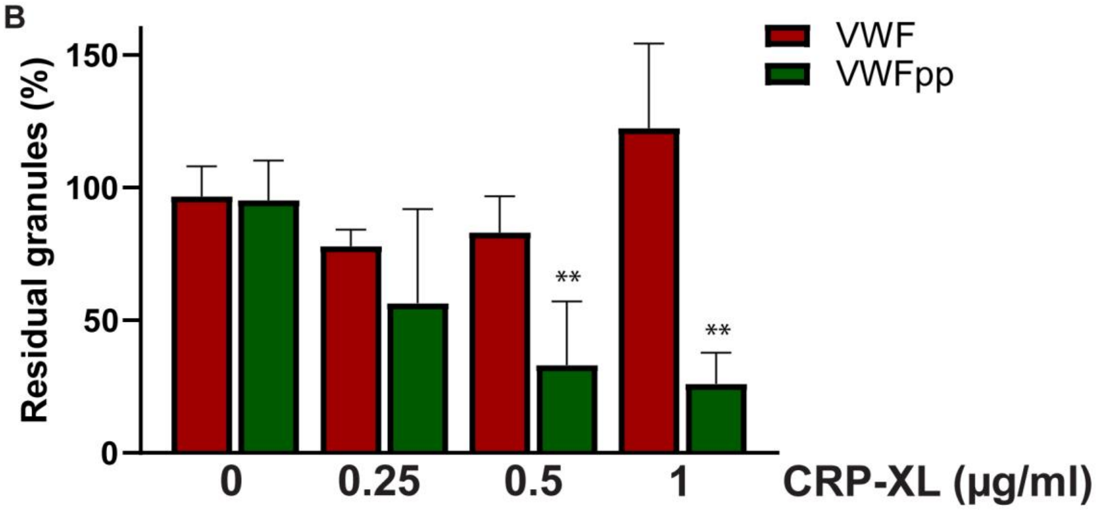

**Supplemental Figure 5: Dose-response release of VWF and VWFpp.** Platelets were stimulated with 0-1 µg/ml CRP-XL and stained for α-tubulin, VWF and VWFpp (A). Representative single plane zoom-in images are shown. Scale bar represents 1 µm. VWF and VWFpp release were assessed by quantification of their residual levels in platelets normalized to resting platelets (B). ** = $p<0.01$, as analyzed by two-way ANOVA. Data presented as mean ± SD.
